# ANIMA: predicting protein-protein interactions across species

**DOI:** 10.64898/2026.09.15.751862

**Authors:** Bruno Florentino, Robson Bonidia, André L. F. de Carvalho, Alexander Schönhuth

**Affiliations:** Institute of Mathematical and Computer Sciences, University of São Paulo, Av. Trabalhador São Carlense, 400, 13566-590, São Paulo, Brazil; Faculty for Technology, Bielefeld University, Universitätsstraße 25, 33615, North Rhine-Westphalia, Germany; Department of Computing, Federal University of Technology – Paraná (UTFPR), Av. Alberto Carazzai, 1640, 86300-000, Paraná, Brazil

**Keywords:** protein-protein interaction, interaction prediction, cross-species, animal proteins, artificial intelligence

## Abstract

**Motivation:** Protein–protein interactions (PPIs) underpin a wide range of biological functions in living organisms. Experimental identification of new PPIs is expensive and time-consuming. The experimental bottleneck has implied an imbalance in terms of data availability: while certain species have been screened exhaustively, other species have not been sufficiently examined. An AI driven protocol for PPI prediction that leverages the massive data accumulated for certain species means a decisive boost for so far understudied species.

**Results:** We present ANIMA (Artificial Neural Interaction Model for Animals), an AI supported cross-species PPI prediction model trained on popular species to predict PPIs in under-researched species. Our experiments demonstrate that our model, when trained on 200 diversely selected animal species, can successfully predict PPIs in other species: ANIMA achieves 95.3% accuracy on other animal species, 91.1% on other eukaryotes, and 82.5% on non-eukaryotes. For a more fine-grained evaluation of the model, we stratify performance rates by the evolutionary distance of test to training sets. We also stratify results by a novel, alignment based score (“representation score”) which allows for fine-grained evaluation in terms of its capacity to generalize to unseen interactions. As expected, results demonstrate increasing performance on increasing evolutionary similarity and on increasing identity of interacting proteins, while still showing excellent performance on proteins entirely lacking counterparts in the training set. In comparison with the state of the art, ANIMA demonstrates substantial superiority in terms of performance rates.

**Availability:** ANIMA GitHub Repository

## Introduction

Proteins are the major functional cellular units and perform a wide range of fundamental biological functions. The majority of these functions are associated with protein–protein interactions (PPIs): approximately 80% of proteins exert their functional activity through such interactions (Liu et al., 2025). This explains why PPIs play a central role in key biological processes, including DNA synthesis (Chen et al., 2019b; Yang et al., 2020), gene transcription and translation (Chen et al., 2019b; Yang et al., 2020), cell proliferation (Yang et al., 2020), and immune response (Chen et al., 2019b; Yang et al., 2020).

Consequently, mapping PPI networks, reflecting the interactomes of species, is essential to understand cellular mechanisms, elucidate pathological processes, and identify molecular targets for therapy (Kosoglu et al., 2024). However, the experimental identification of these interactions is a complex, costly, and time-consuming process (Chen et al., 2019a,b; Yang et al., 2020). The general situation explains why machine learning (ML) / artificial intelligence (AI) tools to predict PPIs, thus avoiding time-consuming experimental investigations and accelerating the PPI discovery process, have become popular in general (Charih et al., 2025).

In this, there is another lever one can pull: while PPI networks for certain species have been extensively researched, the interactomes of less popular species have remained insufficiently examined. This bias in terms of species preference was further emphasized by the development of PPI prediction models that focus on inferring missing PPIs from existing ones within species. Examples of such approaches exist for *Homo sapiens* (Yao et al., 2019), *Helicobacter pylori* (Chen et al., 2019b; Zandi et al., 2023), *Saccharomyces cerevisiae* (Chen et al., 2019a,b; Zandi et al., 2023), and *Arabidopsis thaliana* (Zheng et al., 2023; Zhang et al., 2016). As a result, the interactomes of certain species have been near-complete, while the interactomes of others have remained largely unmapped.

This explains why cross-species PPI prediction approaches, leveraging the near-exhaustive knowledge accumulated for extensively studied species to examine underresearched species, have recently gained increasing popularity. Examples of such approaches are DSCRIPT (Sledzieski et al., 2021), Topsy-Turvy (Singh et al., 2022), TUnA (Ko et al., 2024), DeepNano-seq PPI (Deng et al., 2024), PLT-Interact (Liu et al., 2025) and SENSE-PPI (Volzhenin et al., 2024). These approaches fill an important gap based on the realization that PPIs have evolved even across more distant evolutionary taxa. In fact, all of these approaches have already demonstrated decent performance rates.

All of the approaches just mentioned have been investing in the implementation of fairly sophisticated models emerging from recent advances in large language modeling or approaches that are similar in spirit (see ‘State-of-the-Art Tools’ in the ‘Benchmarking’ section below for details). The core insight of our approach is that, when appropriately set up, easy-maintenance resource friendly multilayer perceptrons (MLPs) achieve performance rates that outperform the more sophisticated and demanding approaches by fairly large margins.

In this vein, we propose ANIMA (Artificial Neural Interaction Model for Animals), a model that is based on MLPs of little depth for performing cross-species PPI prediction. We demonstrate how ANIMA, when trained on a selection of well-studied animals, performs on other species at varying evolutionary distances. To analyze results in a sufficiently fine-grained way, we also stratify results according to the evolutionary distance of proteins in test sets from proteins in the (animal based) training set. Beyond the general advantage over prior approaches, we realize in particular that ANIMA’s gains in performance tend to increase on increasing evolutionary distance, which demonstrates its superiority in terms of generalizing to more distant evolutionary taxa.

## Materials and Methods

In this work, we have developed a machine learning model to classify protein pairs as interacting or not. The driving idea of the approach is to learn how proteins interact from species in which such data are abundantly present and predict interactions in other species. As an exemplary use case, we learn how proteins interact from 200 well explored animal species. We then test the performance of our model on 261 animal species, not considered during training, as well as 2 313 non-animal species among which 856 other eukaryotes, and 1 457 noneukaryotes.

We retrieved interaction data from the STRING database version 12 (Szklarczyk et al., 2023), the largest available repository of PPIs (Yoon et al., 2025) (see Supplementary Figure 1, for exact numbers of interactions), which provided us with positives in the sense of truly interacting protein pairs. Further, we followed common random sampling practices for determining truly non-interacting protein pairs, that is, negatives in terms of machine learning terminology. Both training and test sets are composed of positives retrieved from STRING and negatives obtained by approved random sampling techniques.

Figure 1 provides an overview of the entire process, from species selection to classifier training.

**Figure 1.**
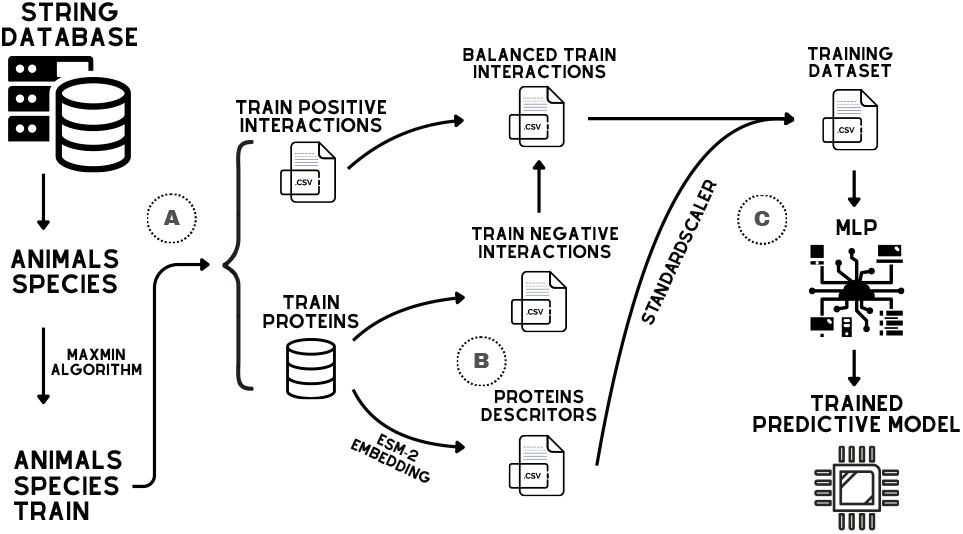
(A) Among the animal species available in the STRING database, a subset is selected using the MaxMin algorithm to compose the training set. Only interactions with a confidence score above 90% and involving proteins with up to 1,000 amino acids are considered. (B) Based on the proteins involved in the positive interactions, negative interaction pairs are generated through random sampling to balance the dataset. In parallel, protein descriptors are extracted using the ESM-2 model with mean pooling. (C) Next, interaction pairs and their corresponding descriptors are concatenated into feature vectors, which are normalized using a Standard Scaler and subsequently used to train an MLP-based classifier.

### Selecting Species for Training

We used the phylogenetic tree provided in the STRING documentation to first identify all 461 animal species. Subsequently, we determined a subset of 200 animal species that is maximally diverse in terms of evolutionary relationships. For the latter, we formalized species selection as a Maximum Diversity Problem (MDP). The MDP is defined to select a subset of *m* elements (here: *m* = 200) from a universe of *n* candidates (here: *n* = 461) such that the sum of pairwise distances between the selected elements is maximized (Martí et al., 2013). The corresponding distances are defined as the minimum number of edges separating species in the phylogenetic tree.

We solve the MDP using the MaxMin algorithm (Gillet, 2011), which starts by initially picking a random species and iteratively selecting the species that is farthest from the ones already chosen. This strategy makes sure that our training set consists of species that are broadly distributed across the phylogeny, promoting high diversity. After having selected 200 animal species this way, whose interactions serve as positives in our training set, we remain with 261 animal species whose interactions serve as the positives in an animal based test set.

### Selecting Positive Interactions and Generating the Negative Sample

To reduce the computational burden, we follow common practice (Ko et al., 2024) and discard all proteins of more than 1 000 amino acids. That is, positives in our training and test sets reflect interaction pairs both proteins of which are at most 1 000 amino acids in length. For the training set, we further filter the interactions and discard all interaction pairs of a confidence score (as provided by STRING) of less than 0.90. Computing the average confidence score of the remaining interactions results yields 0.95, which translates into 95% of the interactions expected to be correct. In summary, we both control for computational expenses and false positives when selecting positives for our training set.

As a result, we have a set of positive interactions along with the primary amino acid sequences of the interacting proteins. To generate negative interactions, that is, protein pairs that do not interact, we randomly sampled pairs of proteins not reported to interact that matched the number of selected positives, thus following a widely adopted strategy (Murakami and Mizuguchi, 2022; Zandi et al., 2023). Importantly, matching the numbers ensures that the training set and subsequently, the test sets are balanced which optimally supports the training and subsequently an unbiased evaluation of results. Although widely approved, of course, this strategy may be affected by the potential mistaking of hitherto undiscovered interactions for negatives. It is important to take such potential labeling errors into account, both when training and testing.

### Feature extraction and training methodology

To ensure smooth machine learning protocols, we further transformed each protein into an equally sized real valued vector (i.e. an *embedding*), using ESM-2 (Lin et al., 2022). In more detail, we opted for the 150-million-parameter version of the provided ESM-2 models, reflecting the optimal choice relative to the available local computational resources. In detail, given an input sequence of *N* amino acids, one obtains an *N ×* 640-matrix to further work with, where each row refers to one of the amino acids in the sequence. A subsequent mean-pooling operation, averaging the *N* different 640-dimensional real valued vectors (each of which is the row for one amino acid) then produces one comprehensive 640-dimensional vector for the entire input protein. When processing pairs of proteins, this amounts to 1280 = 2 *×* 640-dimensional vectors. Adding one extra (binary-valued) dimension to keep track of positive and negative interactions completes the machine learning-adapted representation of non-/interacting protein pairs.

We further normalized the 1280 dimensions across the protein pairs, that is, we subtracted the mean and divided by the standard deviation within each of the 1280 dimensions. Adding the binary-valued label results in 1281-dimensional vectors that one uses as input to the Multilayer Perceptron (MLP) that performs the classification (see (Mussa and Khalifa, 2025) reference where one can look up the definition). We performed hyperparameter optimization exclusively on the training data. Due to computational constraints, we adopted a random search strategy, in which 150 configurations were sampled from the predefined search space (Supplementary Table 1 and 2). For each sampled configuration, we assessed performance using 3-fold cross-validation. Based on the corresponding results, we selected the best-performing hyperparameters and subsequently used them to train the final model on the complete training set.

### Defining Independent Tests

Figure 2 provides an overview of the process for generating the test sets used to evaluate the performance of the predictive model. To evaluate model performance, we first made use of interactions from the 261 animal species not included during training as the primary test set. Guided by the usual phylogenetic trees (see Supplementary Figure 1), we further evaluated our predictive model on nonanimal interactions as additional test sets. In detail, we collected interactions from all species belonging to the Fungi kingdom (663 species), the Plantae supergroup (128 species), and all Eukaryota species not belonging to Animalia, Fungi, or Plantae, commonly referred to as Other Clades (65 species). Furthermore, we considered all interactions from all species belonging to the Archaea domain (457 species) and a set of 1,000 Bacteria species selected using the greedy Max–Min algorithm (Gillet, 2011), the same algorithm that we used to select a sufficiently diverse set of animals for training. According to the phylogenetic tree, these test sets reflect species at gradually increasing evolutionary distances from the training data: First, animal species not seen during training but belonging to the same evolutionary taxon, second, other eukaryotic clades (Fungi, Plantae, and Other Clade eukaryotes) and finally, noneukaryotic domains (Archaea and Bacteria). Our design of test sets facilitates the evaluation of cross-species interaction prediction tools both with respect to within-domain performance and with respect to the capacity of the model to also generalize to diverse, more distant evolutionary groups.

**Figure 2.**
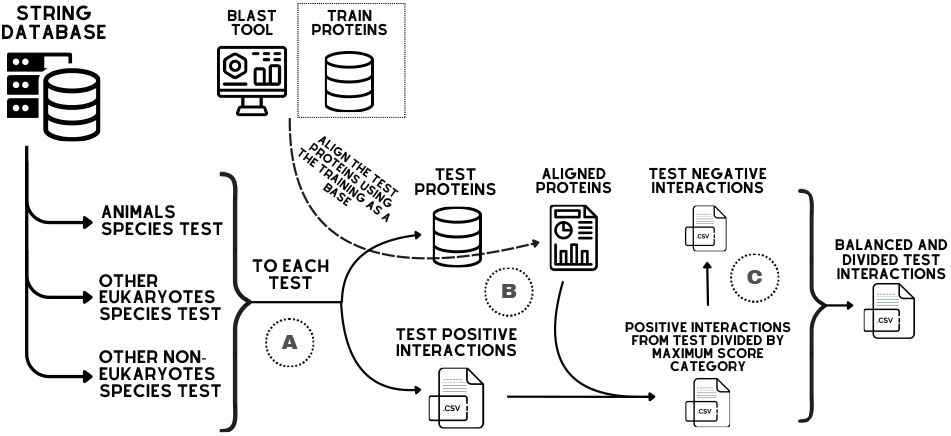
Test workflow. (A) First, all animal species not used during training are collected, together with eukaryotic and non-eukaryotic species from the STRING database. For each group, only interactions with a confidence score of at least 98% and involving proteins of up to 1,000 amino acids are retained. (B) To analyze the relationship between training and test proteins, all proteins from each test set are aligned against the training proteins, and a representation score is computed for each protein. (C) Next, interactions are stratified according to this score, creating subdivisions within the test set. For each subdivision, negative interactions are generated via random sampling to balance the data.

For all these test sets, interactions involving proteins of at most 1,000 amino acids and with a STRING confidence score equal to or greater than 98% were considered. When averaging scores across all test sets, this amounts to 99% of the interactions expected to be correct, or in other words to 1% of incorrectly positively labeled samples. Note that the relative amount of correctly positively labeled samples exceeds the corresponding amount in the training data (training: 95%; test: 98%). In this, we follow common practice (e.g. Sledzieski et al. (2021); Volzhenin et al. (2024)), reflecting that minimizing labeling errors in the test data is essential because labeling errors can decisively blur one’s view on the performance rates of the models.

### Alignment between Training and Test Proteins

In order to support a more fine-grained analysis of performance rates, we seek to stratify interactions in the test sets according to whether test protein pairs have evolutionary counterparts in the training data or not. To this end, we compute the sequence identity, defined for two proteins as the percentage of identical positions among all positions in their mutual alignment. It is common knowledge that high sequence identity results in structurally similar proteins, implying that sequence identity serves as a proxy for evolutionary relatedness (Kanduc, 2012). Guided by these insights, we determine the pair of interacting proteins in the training set that maximally matches interacting test protein pairs in terms of sequence identity, and stratify test protein pairs by the sequence identity of the best matching training protein pair.

We use BLAST (Ye et al., 2006) to calculate the identities between all proteins in the training set and those in the test sets. Because we would like to determine the best matching training proteins for test proteins, we define the training set as the alignment database, run each test protein against that database, and note down the optimal hits. To appropriately stratify the test protein pairs, we define the **representation score** as a new metric for aligned proteins, which ranges from 0% to 100%, and corresponds to the maximum value one obtains from multiplying the aligned fraction of the test protein with the sequence identity of the aligned part. This takes into account that BLAST is a local alignment method, which means that the calculated sequence identity of the aligned protein may refer to an only small part of the alignment (Teufel et al., 2023). Overall, the representation score reflects to what degree the test protein had already been taken into account during training. For example, a representation score of 100% reflects that a test protein had a counterpart in a training pair that matched it at 100% identity and at full length.

For an analysis based on stratifying test protein pairs relative to their representation scores, we first defined 3 different classes for single proteins: *60-100* (*High* or *H*), *40-60* (*Medium* or *M*) and *0-40* (*Low* or *L*), reflecting the representation scores of single proteins. Having stratified single proteins, we further stratified interactions relative to the representation scores of both interacting proteins. This resulted in the classes (*High, High*), (*High, Medium*), (*Medium, Medium*), (*High, Low*), (*Medium, Low*), (*Low, Low*), simply reflecting the representation score classes of the two interacting proteins. Note that these classes can be organized as a lower-triangular matrix. This arrangement enables us to evaluate the capacity of the model to recognize interactions based on the familiarity of the interacting proteins. Of course, one expects the model to perform particularly well for protein pairs belonging to the (*H,H*) class, while the performance of the model on (*L,L*) protein pairs reflects its capacity to generalize to protein pairs that were not considered during training.

### Generating Negative Interactions

We stratify each test set (Supplementary Table 3) relative to the representation score according to the descriptions above. This results in the lower-triangular matrices mentioned above and partitions each test set into 6 subsets. We generate negative samples for each subset using random sampling matching the procedure we used to generate negative samples for the training data set, as described above. For example, when generating negative samples for the (*60–100, 60–100*) = (*H, H*) subset of protein pairs, we randomly select non-interacting protein pairs only from the proteins that make up that subset. Since the amount of positive and negative samples of each subset is balanced, this ensures that we can analyze each subset independently.

## Experiments

We extract protein descriptors for the balanced test sets using the ESM-2 model (Lin et al., 2022), applying mean-pooling to generate a fixed-size embedding (i.e. a real-valued vector of equal length) for each protein. The normalized vectors are fed into the pre-trained MLP model, and, for evaluating the performance of the model, the resulting predicted labels are compared with the true labels.

In the following, a *positive* is a truly interacting protein pair, while a *negative* reflects a non-interacting protein pair, as generated via the negative sampling procedure. Correspondingly, metrics considered during evaluation, such as accuracy, recall, precision and F1-score follow the standard definitions.

Our analysis of the performance of the model focuses on two aspects. First, we examine how performance rates on test sets vary relative to the groups of species that established the test sets. Second, we inspect the relationship between performance rates and the representation scores where the latter give rise to the different parts of the lower-triangular matrices. In other words, we evaluate model performance both relative to evolutionary distance and relative to structural similarity of protein pairs.

### Benchmarking

In order to put our model into perspective with previously achieved performance rates, we compare it with the state-of-the-art for predicting protein-protein interactions, evaluating them on the datasets that we presented above. Namely, we consider D-SCRIPT (Sledzieski et al., 2021), Topsy-Turvy (Singh et al., 2022), TUnA (Ko et al., 2024), DeepNano-seq PPI (Deng et al., 2024), PLT-Interact (Liu et al., 2025) and SENSE-PPI (Volzhenin et al., 2024) for our comparison. Importantly, all these methods include at least one model variant designed for cross-species PPI prediction, rely exclusively on primary protein sequence information for inference, and are publicly available for performing new predictions, which matches the intended application of our approach.

#### State-of-the-Art Tools

In the following, we provide a description of the functional principles of each of the state-of-the-art tools we consider.

**D-SCRIPT** (Sledzieski et al., 2021) encodes protein sequences in structural embeddings using a pre-trained Bi-LSTM (Bepler and Berger, 2019). Pairwise residue interactions are modeled via symmetrized operations and 2D convolutions over a contact map. The resulting features are pooled to produce an interpretable interaction confidence score.

**Topsy-Turvy** (Singh et al., 2022) integrates sequence-based predictions, using D-SCRIPT (Sledzieski et al., 2021) for this purpose, with information derived from the interaction network, incorporating the scores predicted by GLIDE (Devkota et al., 2020) as an additional loss term during training. Although the model leverages the global organization of PPIs only during training, it performs predictions using sequence data exclusively.

**TUnA** (Ko et al., 2024) combines 35M-parameter ESM-2 embeddings, a Transformer encoder to capture intra- and interprotein relationships, and the Spectral-normalized Neural Gaussian Process (SNGP) method to provide reliable uncertainty estimates. The application of spectral normalization to the hidden layers, together with a Gaussian Process-based output layer, enables the model to deliver not only a binary interaction prediction but also a continuous measure of confidence.

**DeepNano-seq-PPI** (Deng et al., 2024) is a cross-species PPI predictor derived from nanobody–antigen interaction modeling. It employs an ensemble of three branches based on different ESM-2 representations, each using a distinct pooling strategy (min, mean, and max) to produce an interaction score. The final prediction combines these scores, with all branches jointly optimized via summed cross-entropy losses. We will refer to the version trained with the 8 million parameter ESM-2 as DeepNano-8, while we will refer to the version trained with the 650 million parameter version as DeepNano-650.

**PLM-Interact** (Liu et al., 2025) uses the 650M-parameter ESM-2 protein language model in a cross-encoder setup, jointly encoding both protein sequences to capture direct residue-level context. A linear layer with ReLU and a sigmoid output then predicts the interaction probability, transferring the structural and evolutionary knowledge learned by ESM-2 to PPI prediction.

**SENSE-PPI** (Volzhenin et al., 2024) encodes protein sequences using 3B-parameter ESM-2 embeddings and processes them with a Siamese GRU to learn independent protein representations. These are combined via a Hadamard product and passed through linear layers to output an interaction probability.

#### Datasets

We evaluate all approaches on the test sets that we composed, and by which one can monitor the models’ performance rates across gradually more distant evolutionary taxa as well as all representation score based classes (which one can visualize as a lower triangular matrix) emerging from the Animalia test species. Following the constraints of the benchmarked methods (Sledzieski et al., 2021; Ko et al., 2024; Deng et al., 2024), we subsampled animal test sets to 5 000 interactions each, the proteins of which were between 50 and 800 amino acids in length; see Supplementary Table 4.

## Results

### Datasets and Experiment Design

Following the methodology summarized in Figure 1A-B and explained in detail in Materials and Methods, we constructed a training dataset comprising 1.8 million positive and 1.8 million negative interactions, encompassing 625,000 proteins distributed across the 200 animal species selected for maximizing diversity. Using this dataset, the predictive model was trained according to the workflow illustrated in Figure 1C with the hyperparameters from Supplementary Table 2. Next, following the methodology summarized in Figure 2A–C, we generated balanced test datasets, as described in Materials and Methods; see Table 1, which reports the number of interactions stratified by the representation scores of the protein pairs.

**Table 1.** Distribution of interaction balanced test pairs across different representation score ranges among the various taxonomic groups, along with the total number of interactions and proteins in each dataset. Labels: *High* (H), *Medium* (M), *Low* (L).

| Test set | H-H | H-M | M-M | H-L | M-L | L-L |
| --- | --- | --- | --- | --- | --- | --- |
| Animals | 624672 | 82122 | 19506 | 29140 | 17124 | 8162 |
| Fungi | 162180 | 230502 | 203970 | 102050 | 337996 | 494732 |
| Plantae | 12364 | 26542 | 27788 | 18836 | 60842 | 86266 |
| Other clades | 22960 | 41306 | 29504 | 15520 | 30666 | 35510 |
| **Eukaryota** | 822176 | 380472 | 280768 | 165564 | 446628 | 624670 |
| **Bacteria** | 46382 | 140450 | 182172 | 150024 | 432508 | 615186 |
| **Archaea** | - | 242 | 15292 | 666 | 46614 | 57164 |

According to Table 1, for animals, the majority of interactions (80%) involve protein pairs in which both proteins have a representation score of at least 60% reflecting the (*High, High*) scenario. In contrast, still considering animals in the (*Low, Low*) category, only a small fraction, approximately 1%, corresponding to 8162 interactions, falls within this range.

Similarly, according to Table 1, the distribution differs for the other eukaryotic groups, *Fungi, Plantae*, and other clades, and other noneukaryotic groups, like the *Bacteria* and *Archaea*. In these additional tests, we observe a low proportion of interactions classified as (High, High), ranging from 0.2% to 11%, with interactions predominantly concentrated among proteins classified as *Low*, and *Medium*.

### Performance on PPIs from Unseen Animals

Using the trained model, we evaluated performance on PPIs involving previously unseen animal proteins to assess generalization to novel cases. On the full animal test set, without stratification by representation score, the model achieved an accuracy of 95%, precision of 93%, and sensitivity of 97%. However, a comprehensive performance analysis requires accounting for the representation score between proteins in the training and test sets. To this end, we evaluated performance across the different representation score based classes defined in Table 1, with the corresponding performance shown in Figure 3 (see Supplementary Table 5 and Supplementary Figure 3 for the extended version in the Supplementary Material).

**Figure 3.**
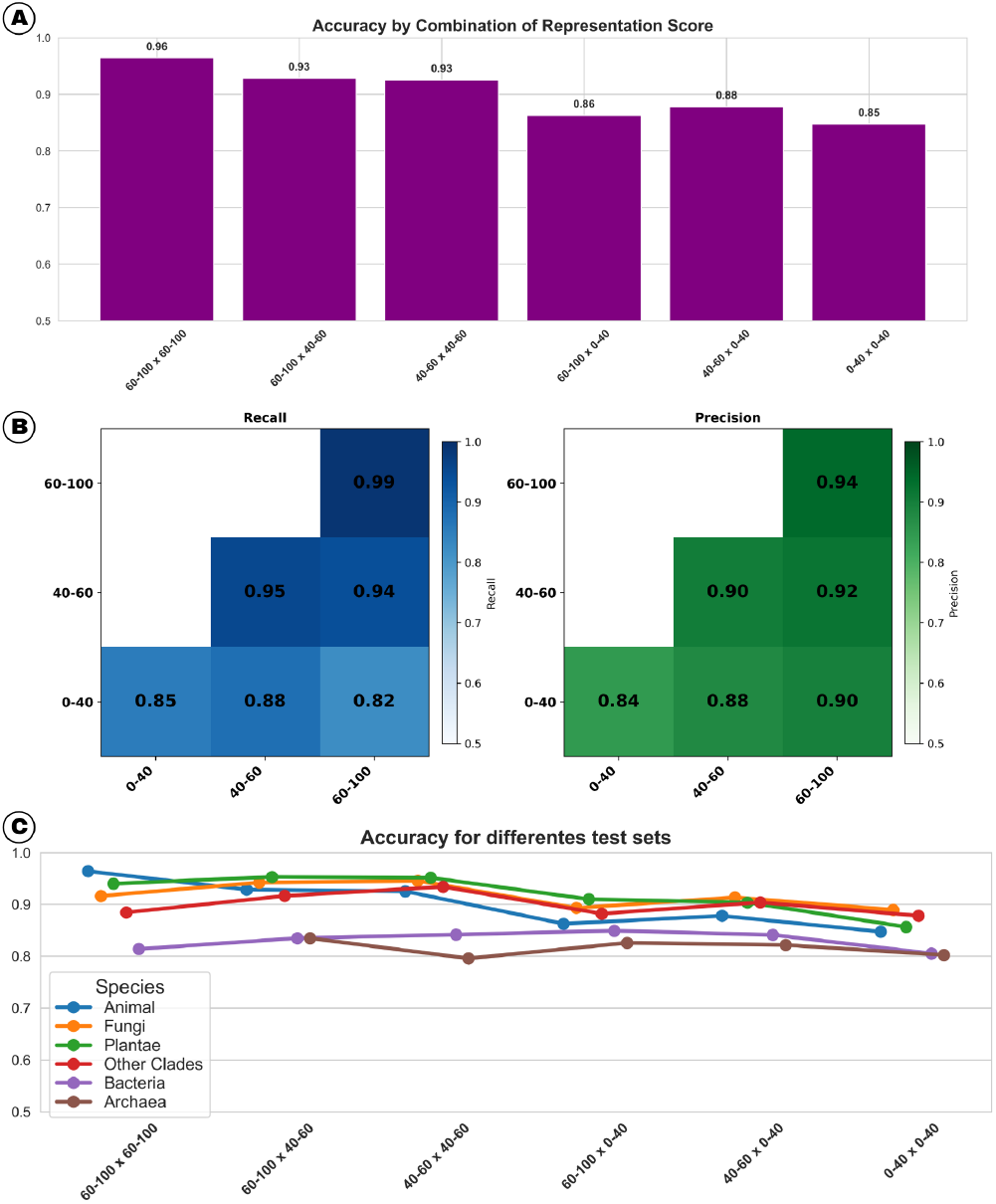
(A) Bar plot of Accuracy for the animal test set, segmented according to the representation score. (B) Triangular performance matrix for the animal test set, segmented according to the representation score. (C) Accuracy performance across different test sets, segmented according to the representation score.

For the animal-based training set, Figure 3A presents the accuracy across different representation score categories. Overall, protein pairs with higher representation scores achieve superior predictive performance, with accuracy ranging from 96% in the (*High, High*) scenario to 85% in the (*Low, Low*) category. In Figure 3B, the precision increases monotonically with higher representation scores. This trend holds both vertically, such that (*High, High*) outperforms (*High, Medium*), which in turn outperforms (*High, Low*), and horizontally, where (*High, Low*) exhibits higher precision than (*Medium, Low*), which in turn surpasses (*Low, Low*). In parallel, sensitivity is highest along the diagonal corresponding to equivalent representation levels (i.e., where *x* = *y*). Along a fixed row, sensitivity increases as one moves upward along the vertical axis, but decreases when moving rightward along the horizontal axis.

### Performance in Different Evolutionary Groups

Expanding the analysis to other test sets listed in Table 1, with predictions made considering the representation score, we have Figure 3C, where the performance in terms of accuracy is displayed. We begin with the (*High, High*) category with 96% F1-score, in which animal proteins exhibit higher performance than the other groups. This is followed by an apparent clustering of fungi (92%), plantae (94%), and other clades (88%) i.e., other eukaryotes, while bacteria (81%) present consistently lower performance than the remaining eukaryotes.

Across the following categories, (*High, Medium*), (*High, Low*), (*Medium, Medium*), (*Medium, Low*), and (*Low, Low*), the performance for animals converges with that for other eukaryotic groups (fungi, plantae, and other clades), which collectively form a cohesive cluster. In contrast, for bacteria and archaea, appear to form a distinct performance group compared to the other eukaryotes, yet consistently exhibit similar performance patterns among themselves as non-eukaryotes.

### Comparing Performance with the State of the Art

In this section, we compare the performance of our model with state-of-the-art approaches; the evaluated datasets are described in Supplementary Table 4. Figure 4 presents the Accuracy results, showing that our model consistently outperforms all competing methods across all scenarios evaluated. Notably, the performance gap widens as the representation score decreases, ranging from approximately 4.5% in Accuracy in the (*High, High*) scenario to 8.7% in in the (*Medium, Low*) scenario, when compared to the second-best model, It also demonstrates better performance than other studies in AUPR, AUROC and F1-score (See Supplementary Tables 6, 7 and 8). These results highlight the robustness of the proposed model for predicting animal PPIs across diverse and increasingly challenging representation settings.

**Figure 4.**
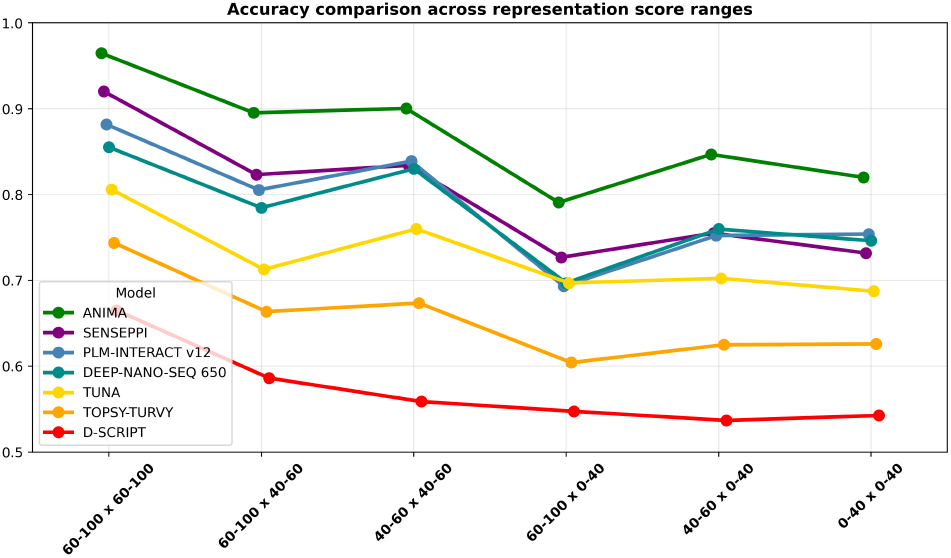
Performance of state-of-the-art tools across different animal datasets.

## Discussion and Final Considerantions

In this study, we developed ANIMA, an AI driven approach for prediction of PPIs across species. We demonstrated that ANIMA when trained on a selection of well-studied animals predicted PPIs in other species at performance rates that were superior over those of the state of the art. From a general perspective, the corresponding improvements yield a considerable boost in terms of leveraging the PPI knowledge gained for heavily researched species to also map the interactomes of less popular species. This avoids the burden of expensive and time-consuming laboratory experiments that are necessary to determine PPIs in a conventional manner.

In order to evaluate ANIMA and its competitors, we invested in carefully designing training and test datasets. While training datasets were required to refer to sufficiently exhaustively examined species and also to span a sufficiently great variety of species in terms of evolutionary diversity, test datasets were supposed to refer to other species at gradually increasing evolutionary distances: in that order, we generated test datasets for other animals, for other eukaryotes, among which Fungi, Plantae and the remaining eukaryotes (‘Other Clades’), and for non-eukaryotes, among which Bacteria and Archaea. As expected, the accuracy achieved by ANIMA in animals not used for training (*≥* 95%) gradually decreased to 91.1% for other eukaryotes to 82.5% for non-eukaryotes.

We also further stratified PPIs in the test datasets by the degree of identity of the participating proteins relative to their counterparts in the training dataset; we refer to the quantity that we introduced and according to which we stratified as the representation score. We realized that the representation score indeed positively correlates with performance rates: the accuracy increases with increasing representation score. Beyond this general insight, we further noticed that ANIMA’s advantages over prior approaches became more striking on smaller representation scores. This provides evidence of the superiority of ANIMA in terms of generalizing to previously unseen proteins over previous approaches.

The key to the success of ANIMA is twofold. First, ANIMA benefit from the phylogenetic tree-based sampling approach that we developed by which to select species for the training dataset, which allowed for sound validation and hence finetuning of models. Second, ANIMA is based on an uncomplicated, and easy-to-train MLP as its core prediction machinery, which renders ANIMA very robust in terms of its usage—for example, when re-training it upon having integrated additional species into the training set.

In summary, we presented an approach to predict PPIs across species that has been shown to be more robust and operate at considerably improved performance rates across the board of evolutionary taxa. Future plans encompass the provision of training datasets for non-animal species and improved definitions of the representation score for even more fine-grained dissection of results into biologically meaningful classes referring to the evolutionary relationship of proteins.

## Data availability

ANIMA is available at: https://github.com/0nurB/ANIMA.

