## Supplemental file for "ANIMA: predicting protein-protein interactions across species"

, , pp. 1–9

### Supplementary material
### ANIMA: cross-species approach for protein-protein interactions prediction

Bruno Florentino,[1,*] Robson Bonidia,[3] André L. F. de Carvalho[1] and Alexander Schönhuth[2]

[1]Institute of Mathematical and Computer Sciences, University of São Paulo, Av. Trabalhador São Carlense, 400, 13566-590, São Paulo, Brazil
[2]Faculty for Technology, Bielefeld University, Universitätsstraße 25, 33615, North Rhine-Westphalia, Germany
[3]Department of Computing, Federal University of Technology – Paraná (UTFPR), Av. Alberto Carazzai, 1640, 86300-000, Paraná, Brazil

#### Abstract

**Availability: ANIMA GitHub Repository**

**Keywords** protein-protein interaction, interaction prediction, cross-species, animal proteins, artificial intelligence

#### String database

##### String score

To use the STRING database version 12 [1, 2] as a source of interactions, it is first necessary to understand how the data are organized. The database encompasses a wide range of species; however, for each species, only interactions among its own proteins are included, with no records of interactions between proteins from different species. Moreover, STRING classifies interactions into confidence levels according to the assigned score:

Following the scoring scheme adopted by the STRING database [1], PPIs were classified into four confidence categories based on their score values: low confidence ($0.15 \leq \text{score} < 0.4$), medium confidence ($0.4 \leq \text{score} < 0.7$), high confidence ($0.7 \leq \text{score} < 0.9$), and highest confidence ($\text{score} \geq 0.9$).

The score provided by STRING is approximately equivalent to the probability that an interaction is true [3]. Thus, when selecting a sample of these interactions, the average score can be interpreted as an approximation of the probability that a randomly chosen interaction within this sample is real.

##### String biodiversity

In parallel, the biodiversity represented in the STRING database [1] encompasses interactions from the three domains of life: *Archaea*, *Bacteria*, and *Eukaryota*. Applying a stringent filter that retains only physical interactions with highest confidence ($\text{score} \geq 0.9$) results in a dataset containing approximately 43 million interactions.

In the Bacteria domain, there are approximately 32.6 million interactions involving 6.6 million distinct proteins distributed across 10,756 species, making it the domain with the largest number of interactions and represented species. In contrast, the Archaea domain comprises about 0.4 million interactions, 0.1 million proteins, and 457 species, representing the domain with the fewest interactions. Specifically, within the Eukaryota domain, approximately 9.2 million interactions are recorded, involving 2.8 million distinct proteins from 1,322 species. The largest species groupings in STRING can be visualized in Figure 1.

A phylogenetic tree is a graphical representation that describes the evolutionary relationships among different species [4]. Each node in the tree represents a common ancestor, while the branches indicate the evolutionary lineages that have diverged over time. This structure can be used to understand the relationships between species: closely related species share a more recent common ancestor than distantly related ones [4]. Thus, it is possible to define groups of species that are evolutionarily closer or more distant.

According to the phylogenetic tree provided by STRING [1], eukaryotic species can be grouped into three main groups: Opisthokonta supergroup [5], Plantae supergroups [5], and Other clades [5], which include representatives from multiple groups such as Stramenopiles, Amoebozoa, and Alveolata. Within these groups, Opisthokonta comprises animals and fungi [6, 5], with 461 and 663 species in the STRING database, respectively. Plantae includes red algae, green algae, and plants [7, 5], totaling 128 species. Finally, the Other clades group encompasses all remaining species that do not belong to the previous supergroups, such as amoebae and protozoa, with 65 species represented in the database.

**Received:** Date Month Year. **Revised received:** Date Month Year. **Accepted:** Date Month Year
© The Author(s) . Copyright and licence statements to be updated by the publisher during production.

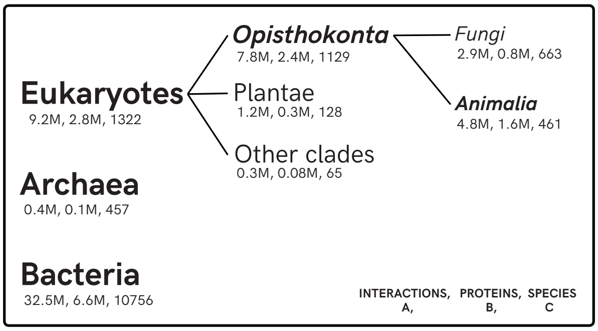


**Figure 1** Major taxonomic groups in the STRING phylogenetic tree, showing the number of PPIs, proteins, and species in each group, considering only physical interactions with an interaction probability of 90% or higher.

#### Hyperparameters

Table 1 summarizes the hyperparameter space explored for the Multi-Layer Perceptron (MLP) during training in order to identify the configuration that maximizes model performance. Subsequently, the selected hyperparameters are reported in Table 2.

**Table 1** Hyperparameter search space used for training the MLP classifier. The model was optimized using the Adam optimizer with the Rectified Linear Unit (ReLU) activation function. Hyperparameter selection was performed via 3-fold cross-validation using only the training data, with a fixed random seed of 42.

| Hyperparameter | Values evaluated |
|---|---|
| Hidden layers | $[512, 128]$, $[512, 256, 128]$, $[1024, 512, 128]$, $[1024, 256, 128, 16]$, $[1024, 512, 256, 128]$ |
| Dropout rate | 0.15, 0.20, 0.25 |
| Batch normalization | True, False |
| Learning rate | $10^{-4}$, $3 \times 10^{-4}$, $10^{-3}$, $3 \times 10^{-5}$ |
| Weight decay | $10^{-4}$, $10^{-5}$, $10^{-6}$, $10^{-7}$ |
| Batch size | 4096, 8192 |
| Epochs | 50, 100 |
| Early stopping patience | 8 |

**Table 2** Selected hyperparameters.

| Hyperparameter | Value |
|---|---|
| Hidden layers | [1024, 512, 128] |
| Dropout | 0.25 |
| Batch normalization | False |
| Learning rate | $3 \times 10^{-4}$ |
| Weight decay | $1 \times 10^{-7}$ |
| Batch size | 8192 |
| Epochs | 100 |
| Early stopping patience | 8 |

#### Test species sets

Table 3 reports, for each test set (Animalia, Fungi, Plantae, Other clades, Bacteria, and Archaea), the number of species used for PPI extraction to construct each dataset.

**Table 3** Number of species across taxonomic groups for the different datasets generated to evaluate model performance, along with their phylogenetic relationship to the Animalia kingdom.

| Test set | Species | Evolutionary relationship |
|---|---|---|
| Animalia | 261 | Same kingdom |
| Fungi | 663 | Same supergroup |
| Plantae | 128 | Different supergroup |
| Other clades | 65 | Different supergroup |
| Bacteria | 1,000 | Different domain |
| Archaea | 457 | Different domain |

#### Datasets for Comparison

Table 4 summarizes the animal PPI datasets used to compare the performance of our model with state-of-the-art approaches, including D-SCRIPT [8], Topsy-Turvy [9], TUnA [10], DeepNano-seq-PPI [11], PLT-Interact [12], and SENSE-PPI [13].

**Table 4** Number of interactions per animal comparation dataset, grouped by representation score categories. The datasets were balanced to contain an equal number of positive and negative PPIs. Only proteins with 50-800 amino acids.

| Category | Number of interactions |
|---|---|
| 60-100, 60-100 | 5000 |
| 60-100, 40-60 | 5000 |
| 60-100, 0-40 | 5000 |
| 40-60, 40-60 | 5000 |
| 40-60, 0-40 | 5000 |
| 0-40, 0-40 | 5000 |

**Table 5** Distribution of interaction balanced test pairs across different representation score ranges among the various taxonomic groups, along with the total number of interactions and proteins in each dataset. **New labels only to this table**: *80-100* (*very High* or *vH*), *60-80* (*High*), *40-60* (*Medium* or *M*), *20-40* (*Low* or *L*), *0-20* (*very Low* or *vL*).

| Test set | vH–vH | vH–H | H–H | vH–M | H–M | M–M | vH–L | H–L | M–L | L–L | vH–vL | H–vL | M–vL | L–vL | vL–vL |
|---|---|---|---|---|---|---|---|---|---|---|---|---|---|---|---|
| Animals | 438214 | 137028 | 49430 | 39872 | 42250 | 19506 | 12652 | 12274 | 14934 | 5674 | 2100 | 2114 | 2190 | 2022 | 466 |
| Fungi | 866 | 13950 | 147364 | 9318 | 221184 | 203970 | 2666 | 77236 | 278092 | 255428 | 1442 | 20706 | 59904 | 174296 | 65008 |
| Plantae | 54 | 1170 | 11140 | 1644 | 24898 | 27788 | 916 | 13204 | 48844 | 49518 | 814 | 3902 | 11998 | 25296 | 11452 |
| Other clades | 966 | 4240 | 17754 | 3542 | 37764 | 29504 | 1096 | 11136 | 25532 | 20058 | 364 | 2924 | 5134 | 11314 | 4138 |
| **Eukaryota** | 440100 | 156388 | 225688 | 54376 | 326096 | 280768 | 17330 | 113850 | 367402 | 330678 | 4720 | 29624 | 79226 | 185928 | 81064 |
| **Bacteria** | 2080 | 10336 | 33966 | 12156 | 128294 | 182172 | 10884 | 109824 | 347580 | 213578 | 2736 | 26580 | 84928 | 175024 | 226584 |
| **Archaea** | 0 | 0 | 0 | 0 | 242 | 15292 | 0 | 576 | 43170 | 42422 | 0 | 90 | 3444 | 10230 | 4512 |

#### Overall accuracy

Figure 2 shows the model's accuracy for each of the test sets.

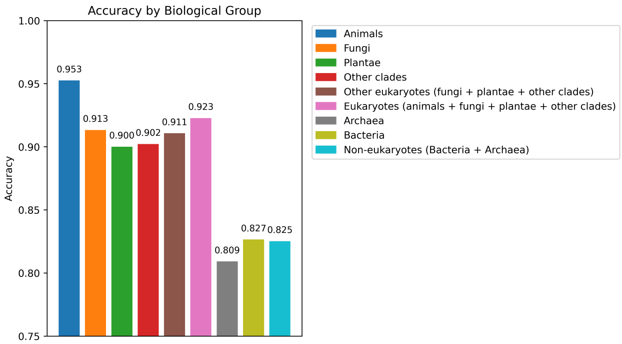


**Figure 2** Accuracy of different test sets.

#### Expanded representation score

We provide the extended version of the figures with the representation score divided into equal 20% intervals. In this new partitioning, the sizes of the datasets are reported in Table 5, where some categories contain a limited number of interactions. The corresponding performance results are shown in Figure 3.

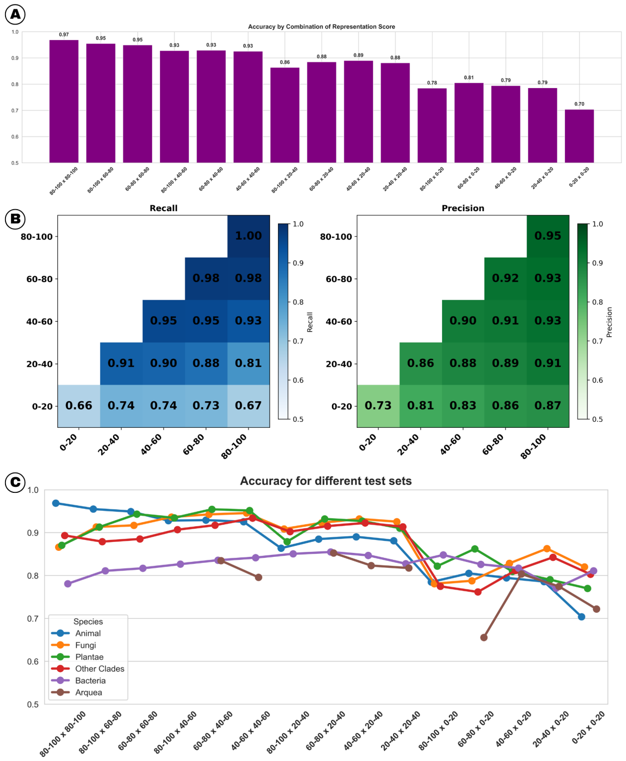


**Figure 3** Extended version with representation score divided into 20% increments. (A) Bar plot of Accuracy for the animal test set, segmented according to the representation score. (B) Triangular performance matrix for the animal test set, segmented according to the representation score. (C) Accuracy performance across different test sets, segmented according to the representation score.

#### Representation Score Histograms

Figures 4, 5, 9, 7, and 8 present the histograms of the representation scores. These scores were obtained by aligning each protein from the test sets with the proteins used for training and computing the corresponding representation metric.

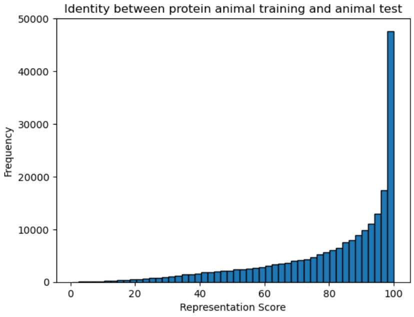


**Figure 4** Distribution of representation scores for proteins from the *Animalia* test set when aligned with the animal training proteins.

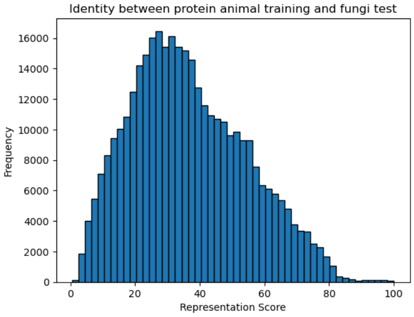


**Figure 5** Distribution of representation scores for proteins from the *Fungi* test set when aligned with the animal training proteins.

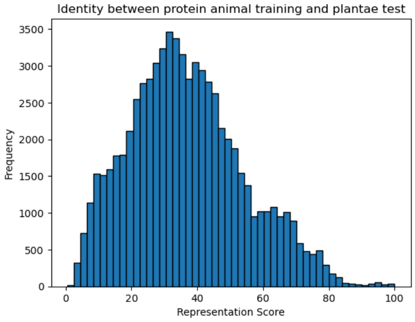


**Figure 6** Distribution of representation scores for proteins from the *Plantae* test set when aligned with the animal training proteins.

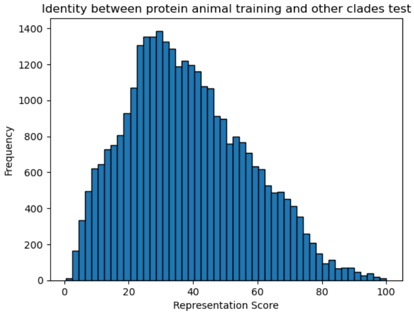


**Figure 7** Distribution of representation scores for proteins from other *Eukaryota* clades, excluding *Opisthokonta* and *Plantae*, when aligned with the animal training proteins.

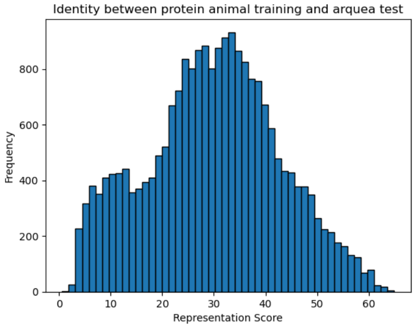


**Figure 8** Distribution of representation scores for proteins from the *Archaea* test set when aligned with the animal training proteins.

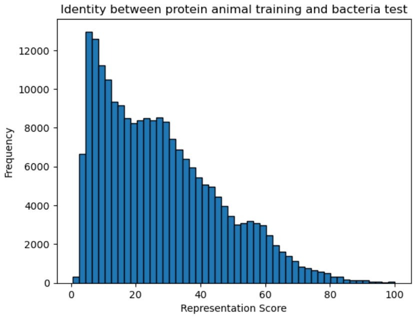


**Figure 9** Distribution of representation scores for proteins from the *Bacteria* test set when aligned with the animal training proteins.

#### Performance on the Test Sets

Figures 10, 11, 12, 13, 15, and 14 present the performance obtained by applying the model proposed in this study to the test datasets described in Table 1 of the main text.

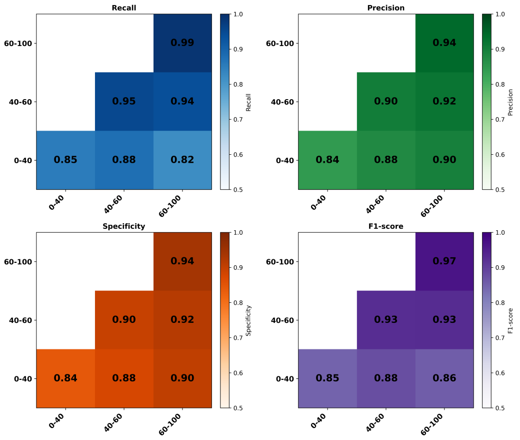


**Figure 10** Triangular performance matrix for the *Animalia* test set.

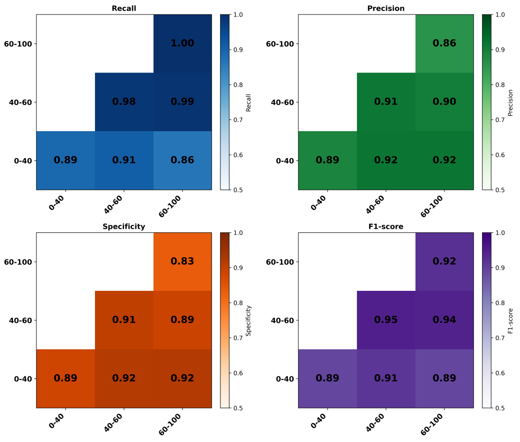


**Figure 11** Triangular performance matrix for the *Fungi* test set.

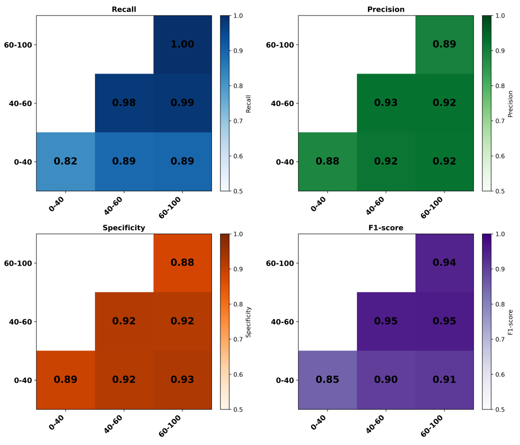


**Figure 12** Triangular performance matrix for the *Plantae* test set.

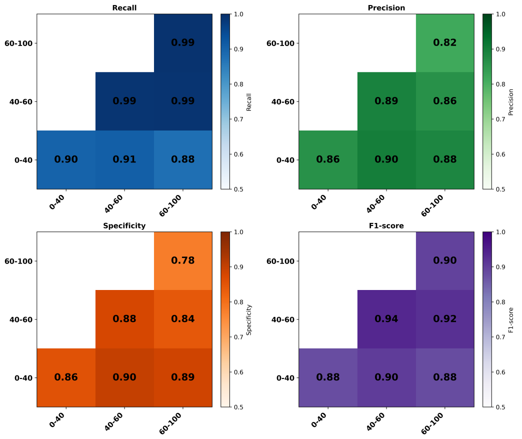


**Figure 13** Triangular performance matrix for other *Eukaryota* clades, excluding *Opisthokonta* and *Plantae*.

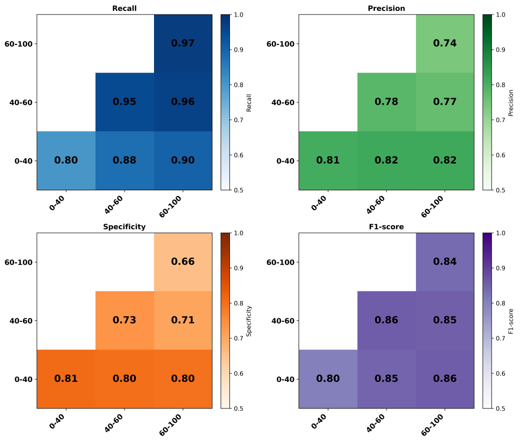


**Figure 14** Triangular performance matrix for the *Bacteria* test set.

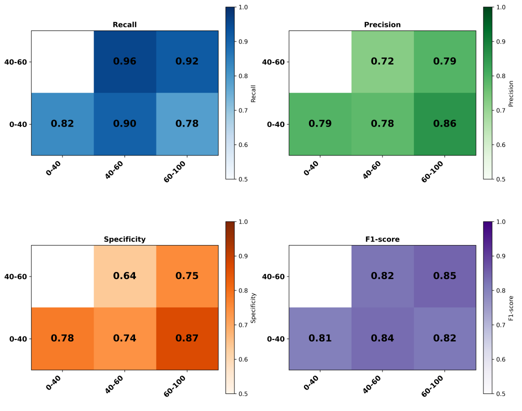


**Figure 15** Triangular performance matrix for the *Archaea* test set.

#### Performance of other state-of-the-art studies

Performance between the various state-of-the-art studies, we have the AUPR curves (Figure 16), AUROC curves (Figure 17) and F1-score (Figure 18).

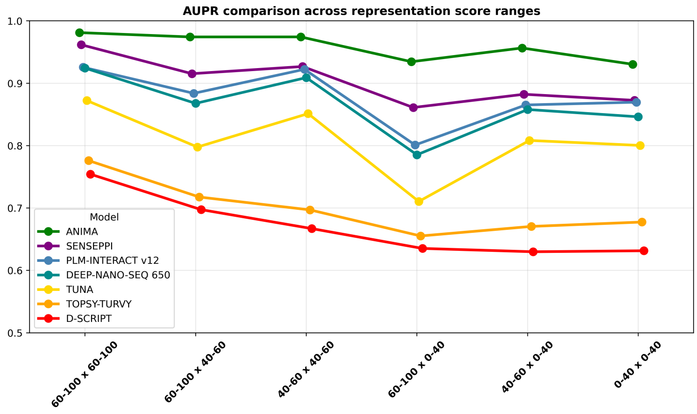


**Figure 16** AUPR performance of state-of-the-art tools across different animal datasets

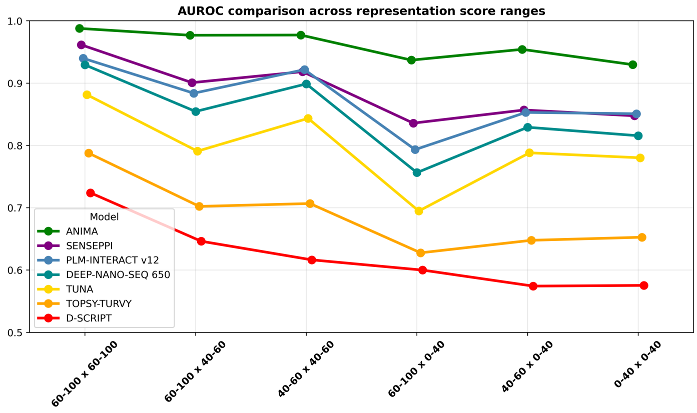


**Figure 17** AUROC performance of state-of-the-art tools across different animal datasets

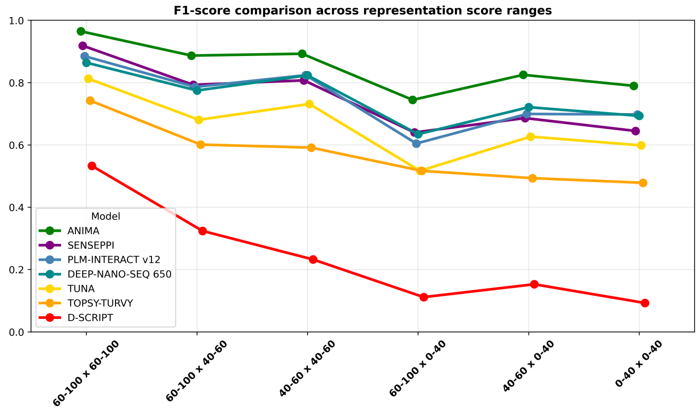


**Figure 18** F1-score performance of state-of-the-art tools across different animal datasets

| Method | H–H | H–M | M–M | H–L | M–L | L–L |
|---|---|---|---|---|---|---|
| Our | 0.9811 | 0.9742 | 0.9743 | 0.9345 | 0.9565 | 0.9304 |
| SENSE-PPI [13] | 0.9617 | 0.9153 | 0.9268 | 0.8611 | 0.8823 | 0.8726 |
| PLM-Interact-v12 [12] | 0.9256 | 0.8838 | 0.9223 | 0.8010 | 0.8652 | 0.8696 |
| PLM-Interact-v11 [12] | 0.8746 | 0.8079 | 0.8657 | 0.7109 | 0.8228 | 0.8256 |
| DeepNano-seq-PPI-650 [11] | 0.9243 | 0.8677 | 0.9090 | 0.7853 | 0.8579 | 0.8462 |
| DeepNano-seq-PPI-8 [11] | 0.8347 | 0.7276 | 0.6375 | 0.7577 | 0.7670 | 0.7396 |
| Tuna [10] | 0.8723 | 0.7977 | 0.8513 | 0.7105 | 0.8083 | 0.8003 |
| Topsy-Turvy [9] | 0.7760 | 0.7176 | 0.6970 | 0.6552 | 0.6705 | 0.6775 |
| D-SCRIPT [8] | 0.7542 | 0.6974 | 0.6670 | 0.6352 | 0.6299 | 0.6315 |

**Table 6** AUPR comparison across representation score categories. H, M, L and denote high, medium and low protein representation levels, respectively.

| Method | H–H | H–M | M–M | H–L | M–L | L–L |
|---|---|---|---|---|---|---|
| Our | 0.9878 | 0.9768 | 0.9772 | 0.9370 | 0.9542 | 0.9297 |
| SENSE-PPI [13] | 0.9615 | 0.9008 | 0.9184 | 0.8358 | 0.8569 | 0.8476 |
| PLM-Interact-v12 [12] | 0.9399 | 0.8838 | 0.9217 | 0.7933 | 0.8530 | 0.8509 |
| PLM-Interact-v11 [12] | 0.8953 | 0.8068 | 0.8656 | 0.6846 | 0.7979 | 0.8019 |
| DeepNano-seq-PPI-650 [11] | 0.9293 | 0.8545 | 0.8988 | 0.7564 | 0.8292 | 0.8156 |
| DeepNano-seq-PPI-8 [11] | 0.8366 | 0.7254 | 0.6145 | 0.7397 | 0.7647 | 0.7239 |
| Tuna [10] | 0.8815 | 0.7908 | 0.8435 | 0.6948 | 0.7882 | 0.7802 |
| Topsy-Turvy [9] | 0.7877 | 0.7022 | 0.7068 | 0.6277 | 0.6478 | 0.6527 |
| D-SCRIPT [8] | 0.7240 | 0.6463 | 0.6164 | 0.6000 | 0.5743 | 0.5754 |

**Table 7** AUROC comparison across representation score categories. H, M, L and denote high, medium and low protein representation levels, respectively.

| Method | H–H | H–M | M–M | H–L | M–L | L–L |
|---|---|---|---|---|---|---|
| Our | 0.9648 | 0.8869 | 0.8928 | 0.7446 | 0.8250 | 0.7898 |
| SENSE-PPI [13] | 0.9186 | 0.7929 | 0.8074 | 0.6401 | 0.6863 | 0.6449 |
| PLM-Interact-v12 [12] | 0.8850 | 0.7861 | 0.8241 | 0.6046 | 0.6995 | 0.6978 |
| PLM-Interact-v11 [12] | 0.8427 | 0.7188 | 0.7869 | 0.5222 | 0.6770 | 0.6536 |
| DeepNano-seq-PPI-650 [11] | 0.8642 | 0.7748 | 0.8234 | 0.6345 | 0.7212 | 0.6933 |
| DeepNano-seq-PPI-8 [11] | 0.7418 | 0.5305 | 0.3603 | 0.4343 | 0.5255 | 0.4381 |
| Tuna [10] | 0.8130 | 0.6807 | 0.7315 | 0.5169 | 0.6268 | 0.5987 |
| Topsy-Turvy [9] | 0.7425 | 0.6012 | 0.5914 | 0.4729 | 0.4934 | 0.4785 |
| D-SCRIPT [8] | 0.5329 | 0.3241 | 0.2324 | 0.1116 | 0.1529 | 0.0928 |

**Table 8** F1-score comparison across representation score categories. H, M, L and denote high, medium and low protein representation levels, respectively.

| Method | H–H | H–M | M–M | H–L | M–L | L–L |
|---|---|---|---|---|---|---|
| Our | 0.9648 | 0.8952 | 0.9004 | 0.7908 | 0.8468 | 0.8200 |
| SENSE-PPI [13] | 0.9202 | 0.8232 | 0.8340 | 0.7268 | 0.7550 | 0.7318 |
| PLM-Interact-v12 [12] | 0.8818 | 0.8054 | 0.8390 | 0.6932 | 0.7522 | 0.7540 |
| PLM-Interact-v11 [12] | 0.8336 | 0.7470 | 0.8050 | 0.6402 | 0.7366 | 0.7240 |
| DeepNano-seq-PPI-650 [11] | 0.8552 | 0.7846 | 0.8300 | 0.6970 | 0.7598 | 0.7462 |
| DeepNano-seq-PPI-8 [11] | 0.7550 | 0.6414 | 0.5760 | 0.6196 | 0.6522 | 0.6204 |
| Tuna [10] | 0.8060 | 0.7128 | 0.7600 | 0.6970 | 0.7024 | 0.6874 |
| Topsy-Turvy [9] | 0.7438 | 0.6636 | 0.6736 | 0.6042 | 0.6250 | 0.6260 |
| D-SCRIPT [8] | 0.6652 | 0.5862 | 0.5588 | 0.5474 | 0.5368 | 0.5426 |

**Table 9** Accuracy comparison across representation score categories. H, M, L and denote high, medium and low protein representation levels, respectively.
